# Isotope analysis of birds’ eye lens provides early-life information

**DOI:** 10.1101/2024.08.28.610164

**Authors:** Emi Arai Hasegawa, Jun Matsubayashi, Ichiro Tayasu, Tatsuhiko Goto, Haruka Inoue, Axel G. Rossberg, Chikage Yoshimizu, Masaru Hasegawa, Takumi Akasaka

## Abstract

Early-life environment has a long-lasting effect on later life, though its estimation is often prevented in the wild because of a lack of available methods. Recently, isotope analysis of eye lenses has attracted considerable interest as a means to reconstruct the environmental conditions experienced by animals during the developmental period. This analysis has mostly been confined to fish for practical reasons and remains to be resolved for application to other animals. In this study, we broadened its applicability by developing a novel approach and verifying its usability for the reconstruction of early-life environments. We performed a feeding experiment using Japanese quail (*Coturnix japonica*), in which we administered two diets: one composed mainly of C3 plants (low *δ*^13^C and high *δ*^15^N) and the other of C4 plants (high *δ*^13^C and low *δ*^15^N). Quails in the control group were continuously fed a C3-based diet from hatching until 200 days old, whereas those in the treatment groups (T10, T15, T20, and T40) were switched from the C3 to the C4-based diet at 10, 15, 20, and 40 days after hatching, respectively. We found that the *δ*^13^C in the eye lenses of the treatment groups decreased from the center layer to the middle layer of the lens and then increased toward the outer layer, thus reflecting the diet change. In contrast, those of the control group exhibited a decreasing trend and equilibrated at the middle layer of the eye lens, with no increase thereafter. This novel approach revealed the postnatal feeding histories of the diet-shift experiment. The high *δ*^13^C values observed in the center of the eye lenses would reflect the prenatal feeding environment, i.e., the C4-based diet consumed by their mothers, which is further reinforced by higher *δ*^15^N values at this position due to the consumption of egg yolk-derived nutrition. These results indicate that the avian eye lens can be used as an “isotopic chronicle,” which is a useful tool for reconstructing chronological isotopic information about their early-life history.

## 1 Introduction

Early-life environment has a long-lasting effect on the later life of animals and is known as the “silver spoon hypothesis” (Monaghan 2008). Food availability, habitat quality, or physiological stress during early life can affect future development, survival, and reproduction (Carlisle et al. 2015; Hayward et al. 2013; Hamel et al. 2009; Nussey et al. 2007). Understanding the early-life environments of organisms is important in pure science to clarify the exact causality of phenotypic development as well as in applied science for the conservation of declining species (Sergio et al. 2022). However, early-life information of grown animals is often challenging to obtain because continuous long-term research is required to track the same individual from natal to adult period, and thus a retrospective approach for the reconstruction of early-life environments is required.

Isotope analysis is a widely used technique for investigating the trophic status and migratory origins of organisms across numerous taxa (Hobson and Wassenaar 2018). For example, isotope analysis plays a pivotal role in elucidating the ecological dynamics of long-distance migrants, such as birds, because of the challenges associated with tracking their migratory patterns and dietary preferences (Inger and Bearhop 2008; Tzadik et al. 2017). Inert integumentary tissues, such as bird feathers, have often been used as exemplary tissues for retrospective isotope analysis (Hobson and Clark 1992; Wolf et al. 2013), as they provide information on their development. In particular, isotope analysis of feathers has played an important role in avian ecology by explaining migratory patterns with a relatively small cost when compared to other methodologies, such as geo-loggers and GPS tracking (Hobson et al. 2012; Guiterrez-Exposito et al. 2015).

However, feathers and other inert integumentary tissues are not flawless because information may be lost after their renewal, i.e., after molting (Carvalho et al. 2022). Therefore, these analyses cannot provide permanent information about early-life environments, such as their birthplaces. Early-life information, which is particularly important, needs to be collected by other, more permanent inert tissues.

One tissue that can potentially preserve the environmental conditions experienced during early life is the eye lens, as eye lenses are composed of metabolically inert proteins that are sequentially deposited (Vecchio and Peebles 2020). Previous studies considering fish eyes indicated that eye lenses can be a valuable tool for reconstructing past environments. However, unlike fish and mammals, avian eye lenses are challenging to analyze because of their softness and water solubility (Simpson et al. 1994); therefore, there is a need to develop a method for avian eye lens analysis.

Avian lens maturation follows a pattern that is common to vertebrates, with new fibers continuously emerging from the equatorial epithelial cells (Sivak 2004). The avian eye lens develops during embryonic stages, and it is postulated that only the outermost lens harbors post-embryonic information (Peebles and Hollander 2020). In addition, during the embryonic stage, synthesized proteins are transported to the aged and metabolically inactive cells at the center of the lens, referred to as the organelle-free zone (OFZ) (Shestopalov and Bassnett 2000), which can be challenging to clarify via isotope analysis of the early-life environment. On the other hand, unlike fish, the avian eye lens does not grow larger for more than 50 days, and also does not undergo compaction at the center of the lens (Augusteyn 2008). Thus, once we can develop a protocol to analyze avian eye lenses, a relatively stable lens size would make a useful tool to reconstruct the developmental environment. By establishing a diet-shift experiment in early life, we can assess whether the eye lens reveals the diet history and, if any, develop a formula to reconstruct the history of the early-life environment from the isotopic information while considering the substantial mixing of the isotopes in the radial direction (i.e., diffusion by the above-mentioned biological transport process and sample preparation process) (Passey and Cerling 2002).

The purpose of this study is to determine whether the environment experienced during the early-life stage can be reconstructed from an avian eye lens. To achieve this, we performed a feeding experiment on captive Japanese quails (*Coturnix japonica*) for 200 days, using diets derived from C3 and C4 plants. After hatching, chicks received a C3-based diet (hereafter C3 food). The C3 food of the control group was maintained throughout the experiment, while the treatment groups switched to a C4-based diet (hereafter C4 food) at specific points (Fig. 1). Following the feeding period, we analyzed a time series of the isotopic ratios in both the eye lens and primary feathers. Based on the data, we aim to (1) test whether birds’ lenses record the chronological isotopic information and (2) determine the duration of such information retention. Furthermore, we investigate the correspondence between the position in the eye lens and the age at which the information is deposited by developing a formula.

**Figure 1.**
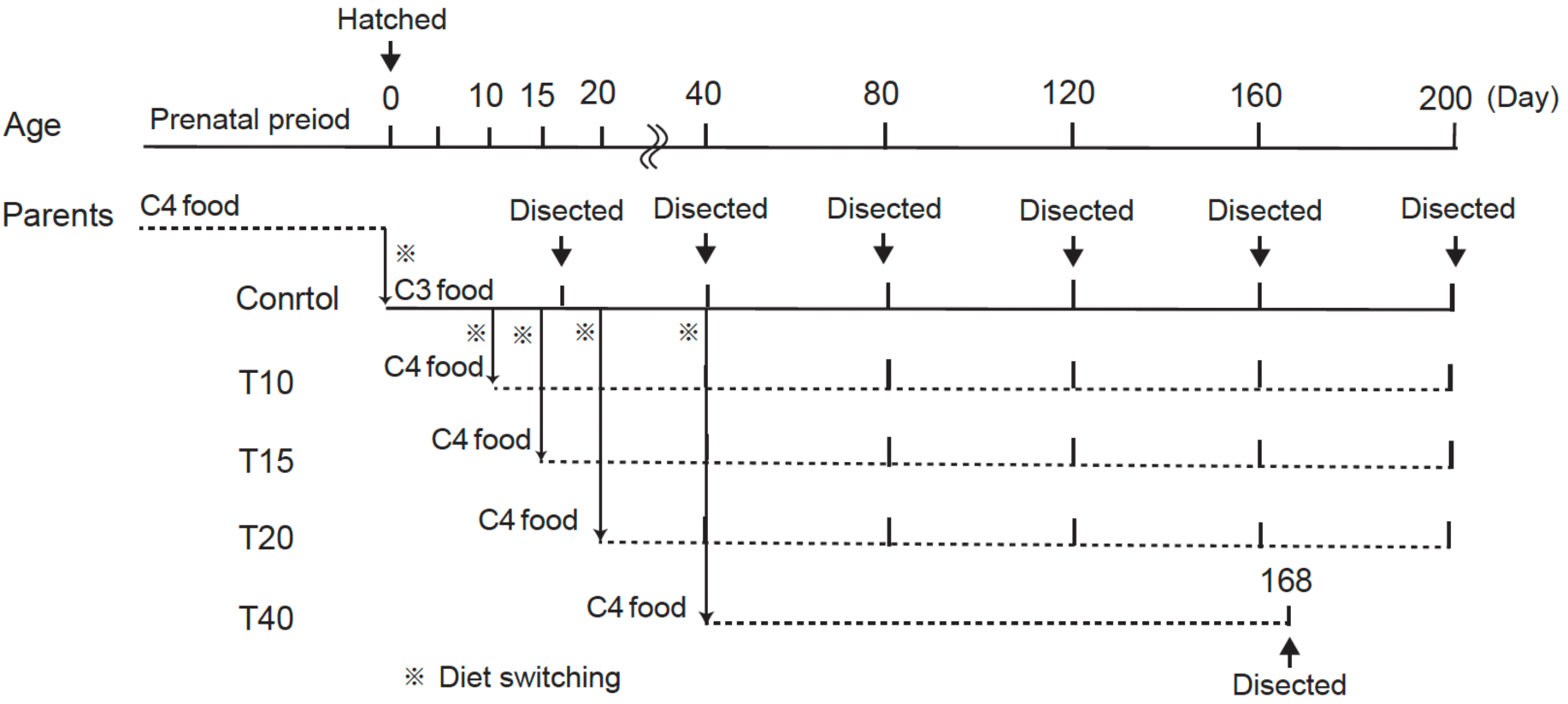
Diet-shift experiment design. Japanese quail eggs were collected from parent birds that were raised on C4 food. The hatched chicks were fed C3 food during the first 10 days, and they were then divided into control group (*n* = 6) and treatment (total *n* = 25) groups. The control group continued on the C3 food until the end of the experiment (i.e., dissection date at days 17, 40, 80, 120, 160, and 200). The treatment group switched to the C4 food at specific time points: 10 days (T10, *n* = 6), 15 days (T15, *n* = 10), 20 days (T20, *n* = 7), and 40 days (T40, *n* = 2) after the start of rearing. The final rearing period (i.e., dissection date) for the treatment groups included time points at 40, 80, 120, 160, 168, and 200 days.

## 2 Methods/Experimental

### 2.1 Diet-shift experiments

For the C3 food (Kusanagi Farm Limited Company, Obihiro, Japan), food residues from potato peels, confectionery waste, cotton and pumpkin seeds, sake lees, and other sources were utilized. This mixture was fermented with lactic acid bacteria and combined with wheat, soybeans, yams, scallops, rice bran, starch flour, fishmeal, and beet cake (Mori et al. 2020). In contrast, the C4 food employed a commercially available mixed feed (Runkeeper, Marubeni Nisshin Feed Co., Ltd., Japan) (Goto et al. 2023), containing imported corn and additional components (Mori et al. 2020). Feeding trials for both diets were performed at a dedicated facility in the Obihiro University of Agriculture and Veterinary Medicine between June 1 and October 20, 2019.

Japanese quail eggs were collected from parent birds that were raised on the above-mentioned C4 food (Runkeeper) (Goto et al. 2023) and were incubated at 36.5°C until hatching (*n* = 31). Hatched chicks were housed individually and fed C3 food for 10 days. They were then divided into control (*n* = 6) and treatment (*n* = 25) groups. The control group was fed C3 food throughout the trial. The treatment group switched to the C4 food at specific time points: 10 days (T10; *n* = 6); 15 days (T15; *n* = 10); 20 days (T20; *n* = 7); and 40 days (T40; *n* = 2) after the start of rearing. The final rearing period for the treatment groups included time points at 40, 80, 120, 160, 168, and 200 days (Table 1). The control group’s final rearing periods were 17, 40, 80, 120, 160, 168, and 200 days.

### 2.2 Eye lens dehydration and dissection

To perform the continuous dissection of the fragile avian eye lens, which is susceptible to liquefaction at room temperature, we established the following dehydration and drying protocol, that was modified from the method described by Matsubayashi et al. (in preparation):

1. After a minor incision in the sclera with scissors, the frozen eyeballs were immersed in 10 mL of absolute ethanol within a screw-capped plastic tube on ice. This initial cut enabled ethanol access to the eye lens while maintaining its structural integrity.
2. Following immersion for 10 min, the eyeball was extracted, the scleral incision was enlarged, and the eyeball was returned to fresh absolute ethanol. This process was repeated with incisions made along the equatorial plane of the sclera. This step enhanced surface dehydration compared to the initial immersion because of the improved ethanol penetration.
3. Thirty minutes after the initial immersion, the eyeball was removed from the ethanol, and the eye lens was carefully extracted using a spatula, tweezers, and scissors. It was then placed in fresh absolute ethanol on ice for 30 min, followed by 60 min at room temperature. This ensured the through dehydration of the entire eye lens through continued ethanol exposure.
4. For complete internal eye lens desiccation, the eye lenses were dried overnight at 40°C in a dedicated drying oven (DO-600FPA, As ONE, Osaka, Japan). The 40°C drying temperature was selected to adjust to the general body temperature of quails (Underwood et al. 1999). The dried eye lens was then dissected according to the protocol described by Matsubayashi et al. (in preparation):

1. The lens was positioned on a Petri dish, photographed under a dissecting microscope (Leica M125, Leica Microsystems GmbH, Wetzlar, Germany) equipped with a Moticam S3 camera (Shimadzu Rika Corporation, Tokyo, Japan), and its diameter was measured from the circumference.
2. The lens was moistened with a minimal amount of deionized water. Using fine-tipped tweezers, the soft material was carefully collected from across the entire surface and transferred to a new designated container.
3. After scraping off approximately 0.5–200.0 µg of eye lens sample for isotope analysis while preserving the overall lens shape, any remaining water was wiped away, and the used instruments were washed or replaced as required.
4. The above steps 1–3 were repeated until the central core of the eye lens was reached.

Because of the inherent aspherical nature and unevenness of the ethanol-dehydrated quail ocular lenses, the short axis was used as the basis for size determination. This defines the lens size as the shortest diameter at the equator (mm). Ethanol treatment for 2 h should not have a significant effect on the stable isotope analysis of epithelial tissue or amino acids, including the eye lens (Borrow et al. 2008; Chua et al. 2020). The analysis excluded the outermost lens section containing the attached fibrae zonulares (Fitzgerald 2018). This discrepancy between lens size and the outermost analyzed position may be due to this exclusion. Finally, individual lens sections were freeze-dried using a freeze dryer (FDU-2100, EYELA, Tokyo, Japan). We excluded samples of the 160-day-old control quail because of sample degradation.

### 2.3 Feather processing

As the first primary feathers had already completely grown by 40-days old, we only analyzed the first primary (P1) feathers (Paritte and Kelly 2009; Wolf et al. 2013) from 40, 120, and 160-day-old birds (control, T10, T15, and T20). The samples were washed with water, soaked in a 2:1 chloroform: methanol solution for 24h, and then rinsed twice with methanol to remove any surface contaminants and oils. After being air-dried in a fume food for more than 24h (Paritte and Kelly 2009), the samples were moved to a drying oven (DO-600FPA, As ONE, Osaka, Japan) at 60°C for more than 24h. The feather vanes were cut using surgical scissors from the tip to the base, approximately 2–3 mm each but approximately 10 mm for the narrow feather tip, which was sufficient for isotope analysis.

### 2.4 Stable isotope analysis

We performed continuous dissection of the fragile avian eye lenses that are susceptible to liquefaction and placed them along with 0.05–0.16 mg of feathers and lens sections, and 0.65 mg of C3 food, and 0.86 mg of C4 food into tin capsules for the carbon and nitrogen stable isotope analyses. Isotope ratios were measured using a Delta V Advantage mass spectrometer (Thermo Fisher Scientific, Waltham, MA, USA) connected to a Flash EA 2000 elemental analyzer via a Conflo IV interface (Thermo Fisher Scientific). The stable carbon (^13^C/^12^C) and nitrogen (^15^N/^14^N) isotopic compositions of the sample (δ_sample_) were expressed in standard delta (δ) notation according to the following formula (Couplen 2011):

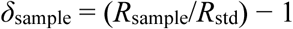

where *R*_sample_ is the corresponding isotope ratio ^13^C/^12^C or ^15^N/ ^14^N for the sample. R_std_ is the defined isotope ratio by international standard scales, where ^13^C/^12^C is the Vienna Pee Dee Belemnite (VPDB) for ^13^C/^12^C and atmospheric nitrogen for ^15^N/^14^N. The isotope ratios of carbon and nitrogen were calibrated against those of the DL-alanine (δ^13^C = –25.36‰, δ^15^N = –2.89‰; CERKU-01), L-alanine (δ^13^C = –19.04‰, δ^15^N = 22.71‰; CERKU-02), and L-threonine (δ^13^C = –9.45‰, δ^15^N = –2.88‰; CERKU-05) laboratory standards, which were calibrated against international standards (Tayasu et al. 2011). The analytical standard deviations (SDs) of the DL-alanine standard (*n* = 53) were 0.09‰ (δ^13^C) and 0.04‰ (δ^15^N); those of the L-alanine standard (*n* = 50) were 0.10‰ (δ^13^C, *n* = 51) and 0.06‰ (δ^15^N, *n* = 56); and those of the threonine standard (*n* = 54) were 0.12‰ (δ^13^C) and 0.07‰ (δ^15^N).

### 2.5 Data analysis

To investigate the correspondence between the position in the eye lens and the age at which the information is deposited, we developed the following model, which has as its starting point a piecewise constant function (PCF), which represents the signature in the eye lens of the switching of diets from the maternal diet (C4 food) to the initial diet (C3 food) and then back to C4 food. This function was convoluted with a normalized Gaussian profile with a standard deviation σ to model the mixing of isotope ratios in the radial direction, which could be caused by contamination of layers with different time frames during the sample preparation or through biological transport process. The convolution evaluates to:

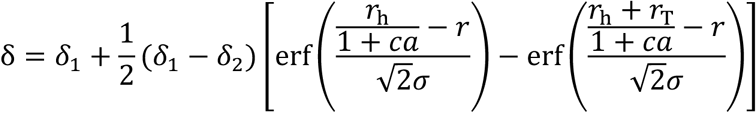

where erf (⋅) denotes the conventional error function; δ_1_ is the asymptotic value for C4 food, i.e., the value where an isotopic equilibrium state is achieved after a change in diet (Ayliffe et al. 2004); δ_2_ is the asymptotic value for C3 food; *a* is the age of quails at the time of dissection; *r*_h_ is the radius corresponding to the time of hatching; *c* is the rate of lens compaction; *r*_T_ is the radius corresponding to the start of the C4 feeding period (*r*_T_ = ∞ for the control); and *r* is the actual distance from the lens center.

We fitted this model to the complete δ^13^C dataset (Supplementary data by minimizing the mean squared difference between δ^13^C observed at radius *r* and the model prediction of δ by adjusting the model parameters σ, δ_1_, δ_2_, *a*, *r*_T_, *c*, and the four independent parameters of *r*_%_ corresponding to the four diet switching times at *T* =10, 15, 20 and 40 days. From this, we estimated the transition lens radius (*r*_h_ + *r*_T_) for each of the dissection date (Supplementary Figure S3). We fitted two model variants, with and without (*c* = 0) lens compaction. In the eye lens of vertebrates, there is a mechanism called compaction, in which the central part of the lens shrinks and does not become larger (Augusteyn 2008), which we account for through the *c* parameter. To date, no studies have examined whether compaction occurs over time in the avian eye lens. Therefore, we constructed models both with and without the compaction parameter (c) and compared them using Akaike information criterion (AIC).

To confirm whether quail lens sizes increased with the dissection date, the relationship between lens size and age (17– 200 day old) was analyzed using a linear model (LM). The isotopic offset between the tissue and the diet (e.g., δ^13^C_lens − diet_) was calculated by subtracting the isotope ratios of the diet from those of the equilibrated tissue.

For the δ^15^N isotope ratio, the asymptotic value was also estimated by following the PCF model:

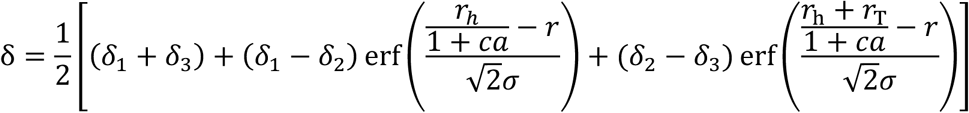

where δ_1_ is the asymptotic value of maternal nutrition, i.e., a high value should be assumed by bioenrichment irrespective of food contents; δ_2_ is the asymptotic value for C3 food; δ_3_ is the asymptotic value for C4 food. The other numerical statements are the same as the above-mentioned PCF model for δ^13^C. All data analyses were performed using the R statistical computing environment version 4.1.2 (R Core Team, 2021).

## 3 Results

### 3.1 Stable isotope ratios of diet and eye lens

The δ^13^C values of the C3 and C4 food were −26.4‰ and −17.1‰, respectively. The PCF model, including compaction, provided a better fit (AIC = 804.5) over the model without controlling for lens compaction (AIC = 837.3, Supplementary Fig. S1). Therefore, the pattern of isotope shift was inferred from the model with compaction (Supplementary Table S1, showing all fitted parameters). In particular, the control quails initially had a high δ^13^C (average ± SD = −19.7‰ ± 1.4‰, with a range of −18.4 to −22.1‰), which decreased with the lens radius (i.e., age), and stabilized at −23.9‰ ± 0.2‰ (Supplementary Table S1). The treatment groups exhibited a peak δ^13^C value in the innermost lens sections, which then dropped and rose again towards the outer sections (Fig. 2). Treatment groups achieved maximum δ^13^C values in the outermost lens sections, which approached the equilibrated δ^13^C value of the C4 food (Supplementary Fig. S1, −17.6‰ ± 0.3‰). Individual variation in δ^13^C existed even among quails with the same rearing conditions (Fig. 2).

**Figure 2.**
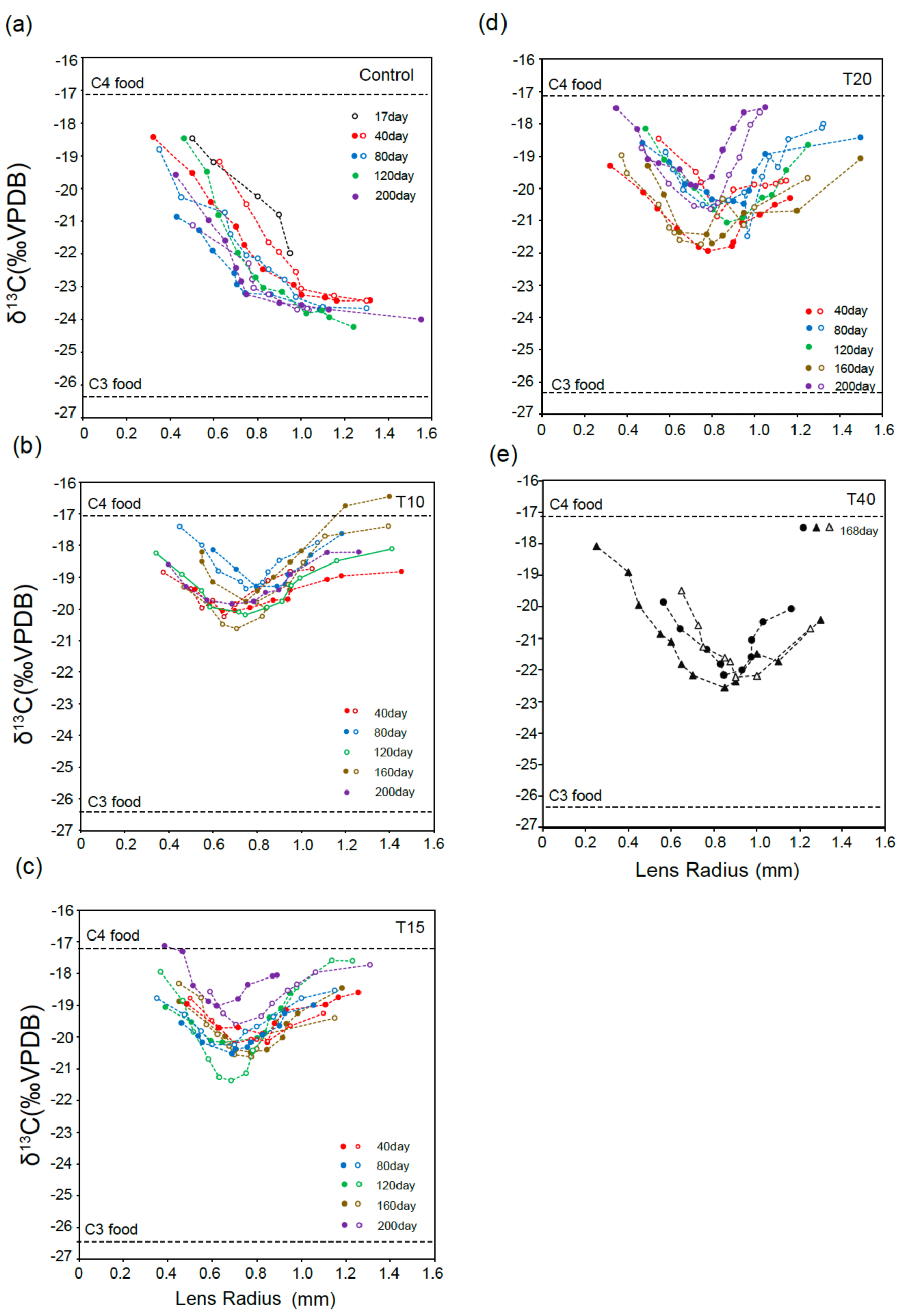
^13^C values of quail eye lenses. (a) Control group (*n* = 5); (b) T10 group (*n* = 6); (c) T15 group (*n* = 10); (d)T20 group (*n* = 7); and (e) T40 group (*n* = 2; 168 days). The lamina from each eye lens (right or left) was included.

The δ^15^N values of the C3 and C4 foods were 4.6‰ and 2.1‰, respectively. The PCF model that included the compaction parameter *c* provided a better fit (AIC = 388.5) compared to the model without *c* (AIC = 416.9, Supplementary Fig. S2). Therefore, the isotope shift pattern was inferred from the model with compaction (Supplementary Table S2, showing all fitted parameters). The δ^15^N values in the center of the eye lens were approximately 5.7‰ ± 0.1‰, which decreased with age and reached equilibrium at 4.5‰ ± 0.1‰ (Supplementary Table S2, Fig. 3(a)). The offset of δ^15^N between the C3 food (4.6 ‰) and eye lens (δ^15^N _lens-diet_) was estimated to be −0.1 ‰. The treatment group also showed a δ^15^N shifts (i.e., decrease) from the center to outer eye lens sections, achieving equilibrium at 3.9‰ ± 0.1‰ (Supplementary Fig. S2, Supplementary Table S2), with individual variations, especially at the innermost sections (Fig. 3). The offset of the δ^15^N between the C4 food (2.1 ‰) and eye lens (δ^15^N _lens-diet,_ 3.9 ‰) was estimated to be 1.8 ‰.

**Figure 3.**
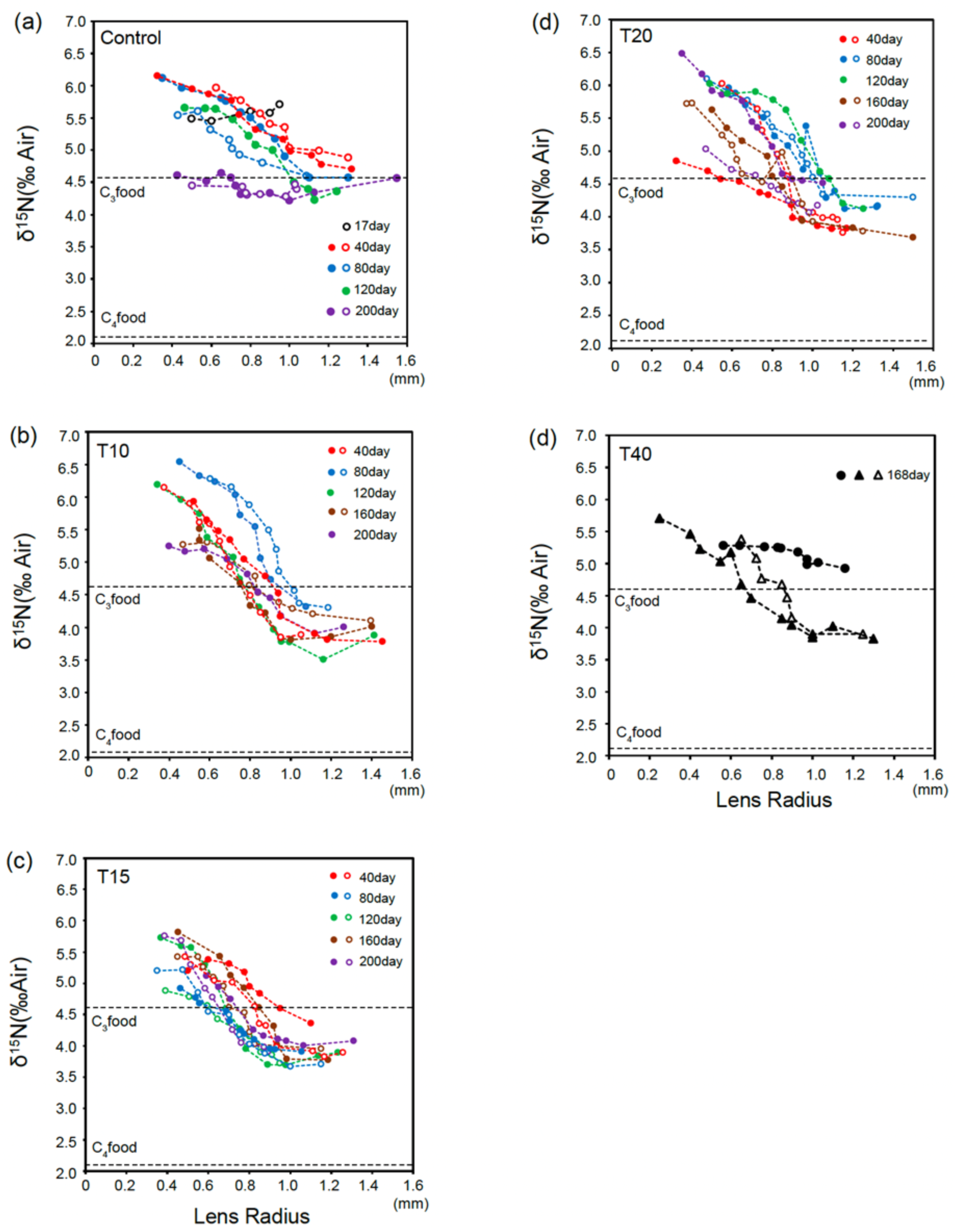
δ^15^N values of quail eye lenses: (a) Control group (*n* = 5); (b) T10 group (*n* = 6); (c) T15 group (*n* = 10); (d) T20 group (*n* = 7); and (e) T40 group (*n* = 2; 168 days). The lens sections from each of eye lens (right or left) were included.

### 3.2 Lens size and compaction

The mean lens size was 1.00 ± 0.07 mm at 17 days (*n* = 2), 1.23 ± 0.13 mm at 40 days (*n* = 8), 1.21 ± 0.15 mm at 80 days (*n* = 8), 1.22 ± 0.15 mm at 120 days (*n* = 6), 1.34 ± 0.11 mm at 160 days (*n* = 7), 1.24 ± 0.07 mm at 168 days (*n* = 3), and 1.10 ± 0.13 mm at 200 days (*n* = 7). There was no significant increase in the eye lens radius with age (from 17 days to 200 days old, *n* = 41, LM: Coef. ± SE = 0.000082 ± 0.0004, t = 0.21, *p* = 0.83). From the PCF models, the lens compaction rate was estimated to be 0.0012 mm per day for the δ^13^C dataset (Supplementary Fig. S3(a), Supplementary Table S1) and 0.0013 mm per day for the δ^15^N dataset (Supplementary Fig. S3(b), Supplementary Table S2).

### 3.3 Stable isotope ratios of primary feathers

We determined the δ^13^C and δ^15^N values of the P1 feathers from quails at 40 days, 120 days, and 160 days (control, T10, T15, and T20 for each). The feather δ^13^C values in the control group remained almost constant throughout the feather section (Fig. 4, mean ± SD = −23.6‰ ± 0.2 ‰, *n* = 3). In the treatment group, all 40-day-old quails exhibited lower δ^13^C values in the tip sections, which increased with growth and reached equilibrium around the δ^13^C value of the C4 food (Fig. 4). The other birds showed almost constant δ^13^C values (Fig. 4, −16.7‰ ± 0.4 ‰).

**Figure 4.**
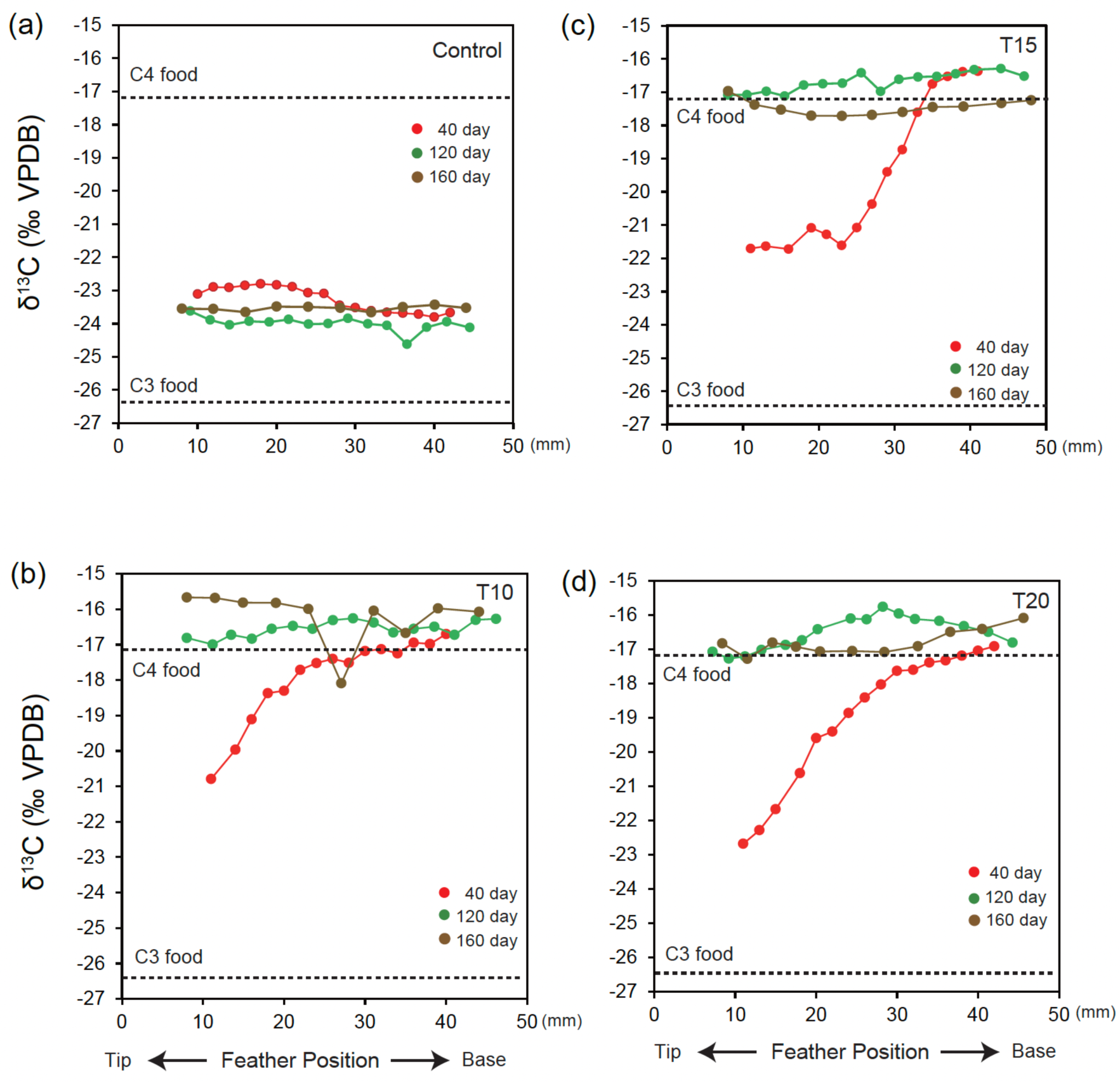
δ^13^C values of quail feather sections. (a) Control group (*n* = 3); (b) T10 group (*n* = 3); (c) T15 group (*n* = 3); and (d) T20 group (*n* = 3).

The feather δ^15^N values of the 120 and 160-day-old quails in the control and treatment groups exhibited individual differences but remained consistent within the individuals (mean ± SD; control group: 3.9‰ ± 0.5‰, treatment groups: 3.4‰ ± 0.5‰, Supplementary Fig. S3). In the two 40-day-old quails, the initial δ^15^N value was high and decreased as they grew, eventually reaching equilibrium around the similar value of the 120 and 160-day-old quails of the same group.

### 3.4 Trophic discrimination factors (TDF) of the quail eye lens and primary feather

We estimated the trophic discrimination factors (TDFs) (i.e., the offset) between each food and tissue. The offset of δ^13^C values between the eye lens (−23.9‰ for the constant value of the control group) and the C3 food (δ^13^C_lens-diet_) was estimated to be 2.5‰ (Supplementary Table S1). The offset of δ^13^C values between the eye lens (−17.6‰ for the constant value of the treatment group) and the C4 food (δ^13^C_lens-diet_) was estimated to be −0.5‰ (Supplementary Table S1). The offset of δ^15^N values between the eye lens and the C3 food (δ^15^N_lens-diet_) was estimated to be −0.1 ‰ (4.5‰, Supplementary Table S1). The offset of δ^15^N values between the eye lens and the C4 food (δ^15^N_lens-diet_) was estimated to be 1.8‰ (3.9‰ for constant value of treatment group, Supplementary Table S1).

The offset of the δ^13^C values between the feather (−23.6‰ for the constant value of the control group, Fig. 4 (a)) and the C3 food (δ^13^C_feather-diet_) was estimated to be 2.8‰ (Supplementary Table S1). The offset of the δ^13^C values between the feather (16.7‰ for the constant value of the treatment group, Fig. 4 (b)−(d)) and the C4 food (δ^13^C_feather-diet_) was estimated to be 0.4‰. The offset of the δ^15^N values between the feather and the C3 food (δ^15^N_feather-diet_) was estimated to be −0.7 ‰ (3.9‰ for the control group, Supplementary Fig. S4 (a)). The offset of δ^15^N values between the feather and the C4 food (δ^13^C_feather-diet_) was estimated to be 1.3‰ (3.4‰ for the constant value of the treatment group, Supplementary Fig. S4 (b)−(d), Supplementary Table S1).

## 4 Discussion

In this study, we found that the early-life environment could be reconstructed using the eye lens from quails. The high δ^13^C values observed in the center of the eye lens for all individuals may reflect the isotope ratios of yolk-derived nutrition, specifically the C4 food fed to their mother (Fig. 2). Subsequently, in the control group of quail that was continuously fed the C3 food, the δ^13^C values in the eye lens transitioned to lower values and equilibrated. In the quails of the treatment groups, the transition to lower C3 food-derived δ^13^C was followed by a shift to higher C4 food-derived values, and this U-shaped shift was preserved in the quails at all switching times (10, 15, 20, and 40 days, Fig. 2). Thus, we confirmed the diet shifts up to 40 days, and hence the quail eye lens should retain isotopic information for at least 40 days after hatching. Furthermore, the decreasing δ^13^C shift from the center to the outer lens was consistent within the control group from 40- to 200-day-old quails (Fig. 2), indicating that the early-life environment is conserved for at least 200 days.

To estimate the timing at which the most recent isotopic information is obtained from the avian eye lens, it is necessary to understand their growth pattern. No significant increase in lens radius was found in individuals after 40 days of age, which is supported by the previous studies in which the avian eye lens did not enlarge after 50 days of age (e.g., house sparrows, *Passer domesticus*, Augusteyn, 2014). Even if lens growth does not occur after its size reaches the asymptote, lens compaction allows for the further preservation of isotopic records (Augusteyn 2008). In our study, the PCF model of both δ^13^C and δ^15^N datasets indicated the presence of lens compaction (Supplementary Fig. S3), but a previous study reported that the absence of lens compaction in chickens (*Gallus domesticus*) has also been observed (Augusteyn 2014). In this study, the longest time to diet shift was 40 days; thus, to determine how long the isotopic record can be stored, it is necessary to conduct experiments with a longer time to diet shift.

We also revealed differences in the isotope ratios between feathers, which have been the most commonly used tissue for isotopic studies in birds (Hobson and Wassenaar 2018), and the eye lens, which we propose as a new target for retrospective isotope analysis. Unlike eye lenses, feathers could not recover embryonic δ^13^C records (Fig. 4), thereby indicating that lenses can recreate the early-life environment, which cannot be obtained from feathers. In addition, in feathers, the transitions from C3 to C4 foods were reflected only in the isotope ratios of quails in the 40-day-old treatment group but not in the 120- and 160-day-old groups, indicating that their early-life environment was lost due to molting. This can be a major obstacle in inferring the isotope ratios (Hobson and Wassenaar 2018). On the other hand, feathers provide isotopic information during feather development, which can be directly linked to information on feather growth. Thus, we showed that the sequential isotope analysis of avian eye lenses provides a new method to complement information on their prenatal phase that is challenging to obtain from feathers.

Although the feeding trial of the current study was designed primarily for δ^13^C, the results of δ^15^N generally supported those of δ^13^C (Fig. 3). The higher δ^15^N values near the center of the eye lens can be explained by the trophic enrichment associated with the consumption of egg yolk-derived nutrients (Hobson1995). When purposely analyzing δ^15^N of the eye lens to understand the trophic ecology via the relationship between animals and their food, this effect of trophic enrichment should be considered. The continuous decline reflecting the diet shift from C3 food to C4 food (Fig. 3) indicated that the avian eye lens also preserves a chronological record of δ^15^N from the prenatal to early postnatal periods. For the δ^15^N of feathers, there was a large variation in the isotope ratios at equilibrium, stored period, and the direction of isotopic ratio among individuals (Supplementary Fig. S4). These irregularities may be linked to feather wear, such as mechanical abrasion and UV light causing a loss of melanin (Johnson 2019), and the source of amino acids, such as endogenous reserves and fed proteins (Cherel et al. 2005), but they do not affect the conclusions obtained for δ^13^C.

The TDFs between the food and tissue in the target organisms are important information when using isotope analysis in ecological studies, which can help predict the isotopic composition of the diets (Hobson and Clark 1992). Because this factor is tissue-dependent, it is worthwhile to estimate the TDFs of eye lenses and feathers to reconstruct the history of diet consumption in a comparable form (Martinez del Rio et al., 2009). The TDFs in quail eye lens and feathers determined in this study (C3 food: δ^13^C_lens-diet_ = 2.5‰, δ^13^C_feather-diet_ = 2.8 ‰, Supplementary Table S1) were similar to those of feathers from a total of 25 bird species (δ^13^C_feather-diet_ = 2.2‰, Swan et al. 2019), but higher than those of whole blood (0.4‰, Swan et al. 2019). The TDFs based on δ^15^N in the current study (C3 food: δ^15^N _lens-diet_ = −0.1‰, δ^13^C_feather-diet_ = −0.7‰, Supplementary Table S1) were estimated to have a lower value than those of previous studies (δ^15^N_feather-diet_ = 4.1‰, Swan et al. 2019, Supplementary Table S1). This lower TDF estimation may partly be related to the difference in the reflection speed of the isotope ratio of the components that are used as a carbon and nitrogen resources via exogenous (mixed feed) and endogenous (stored nutrients) sources depending on their developmental stage (Ayliffe et al. 2004). Furthermore, the mixed feed used in the current study included multiple food sources; therefore, variation in the chemical characteristics (carbon chain length, number of double bonds, degree of unsaturation) between the food components could affect digestibility and energy use in quails (Valentim et al. 2023). These are unavoidable and an intrinsic problem in the TDF estimation using mixed feed. Future research should consider the developmental stages of birds and the turnover rate of their dietary sources (West et al. 2004; Ayliffe et al. 2004).

## 5 Conclusions

In the current study, we developed a methodology for the isotope analysis of avian eye lenses and validated its retrospective use for the reconstruction of early-life environments. Through diet-shift experiment, we were able to reconstruct the early-life history deposited in the avian eye lens and develop a new formula. Our method of dehydration by ethanol and dissection using fine-tipped tweezers is essential to prevent the disintegration of the avian lens. This method can be applied not only to pure science studies but also to applied studies for the reconstruction of the early-life environment for declining species and the implementation of conservation efforts. Although there were large individual differences in the isotope ratios within each treatment group, the control group converged to a single asymptotic value, indicating its reproducibility. Generally, hatchlings (both in precocial and altricial birds) remain in their birthplace, are not expected to disperse within a few weeks, and thus have sufficient time to store their early-life environment within their eye lens, thus making a birds’ eye lens as a valuable tool to reconstruct their early-life environment. Further research is required to improve the accuracy and reproducibility of reconstructing the early-life environments left in the bird eye lens.

## Abbreviations

Air: Atmospheric nitrogen

OFZ: Organelle-Free Zone

PCF: Piecewise Constant Function

TDFs: Trophic Discrimination Factors

VPDB: Vienna Pee Dee Belemnite

## Declarations

### Availability of data and material

The datasets supporting the conclusions of this article are available in the odf.io repository after acceptance.

## Authors’ contributions

JM and EA contributed equally to this work and are co-first authors. JM and TA proposed the topic and conceived and designed the study. TG, and HI performed the feeding trial. JM, EA, and IT developed the method for eye lens dissection and dehydration. EA, CY, JM, and IT performed isotope analysis. AGR developed the formula for data analysis. JM and EA wrote the initial draft of the manuscript. JM, EA, MH, AGR and IT collaborated on the data analysis and its interpretation and the construction of the manuscript. All authors have read and approved the final manuscript.

## Authors’ information

EAH was employed by a grant from JSPS KAKENHI 21H04784 for IT to conduct this research.

## Acknowledgements

Thanks to the laboratory members of the Research Institute for Humanity and Nature for their assistance in our experiments. EA gratefully acknowledges the JSPS KAKENHI 20KK0165 for the opportunity to attend the IsoEcol 2024 held at the University of New Brunswick, Canada.

## Ethics statement

This study was approved by the Experimental Animal Committee of the Obihiro University of Agriculture and Veterinary Medicine (authorization number 19-32).

