## Supplementary Material for "Isotope analysis of birds’ eye lens provides early-life information"

Table S1. All fitted parameters of the piecewise constant function (PCF) model for the $\delta^{13}C$ value.

| Term | Estimate ± SE | t | P-value |
|---|---|---|---|
| $\delta_1$ | −17.6 ± 0.3 | −67.4 | <0.001 |
| $\delta_2$ | −23.9 ± 0.2 | −142.9 | <0.001 |
| $\sigma$ | 0.28 ± 0.02 | 12.7 | <0.001 |
| $c$ | 0.0012 ± 0.0002 | 5.2 | <0.001 |
| $r_{\mathrm{h}}$ | 0.68 ± 0.03 | 24.5 | <0.001 |
| $r_{\mathrm{T10}}$ | 0.26 ± 0.04 | 6.2 | <0.001 |
| $r_{\mathrm{T15}}$ | 0.31 ± 0.04 | 7.0 | <0.001 |
| $r_{\mathrm{T20}}$ | 0.41 ± 0.04 | 9.2 | <0.001 |
| $r_{\mathrm{T40}}$ | 0.70 ± 0.06 | 11.3 | < 0.001 |

$\delta_1$ is the asymptotic value for C4 food; $\delta_2$ is the asymptotic value for C3 food; $\sigma$ is a normal distribution of the width; $r_{\mathrm{h}}$ is the radius corresponding to the time of hatching; $c$ is the rate of lens compaction; and $r_{\mathrm{T}}$ is the radius corresponding to the start of the C4 feeding period. The residual standard error (SE) is 0.7031 at 363 degrees of freedom. The number of interactions to convergence is 13. The achieved convergence tolerance is $5.055e^{-06}$.

Table S2. All fitted parameters of the piecewise constant function (PCF) model for the $\delta^{15}N$ values.

| Term | Estimate ± SE | t | P-value |
|---|---|---|---|
| $\delta_1$ | 5.7 ± 0.09 | 60.9 | <0.001 |
| $\delta_2$ | 4.5 ± 0.08 | 54.5 | <0.001 |
| $\delta_3$ | 3.9 ± 0.08 | 50.0 | <0.001 |
| $\sigma$ | 0.2 ± 0.03 | 5.0 | <0.001 |
| $c$ | 0.0013 ± 0.0003 | 4.9 | <0.001 |
| $r_{\mathrm{h}}$ | 0.86 ± 0.03 | 26.6 | <0.001 |
| $r_{\mathrm{T10}}$ | 0.2 ± 0.07 | 2.8 | <0.01 |
| $r_{\mathrm{T15}}$ | -0.4 ± 0.10 | -3.7 | <0.001 |
| $r_{\mathrm{T20}}$ | 0.3 ± 0.07 | 4.7 | <0.001 |
| $r_{\mathrm{T40}}$ | 0.6 ± 0.18 | 3.4 | <0.001 |

$\delta_1$ is the asymptotic value for maternal nutrition (i.e., the high value should be assumed by bioenrichment irrespective of food content); $\delta_2$ is the asymptotic value for C3 food; $\delta_3$ is the asymptotic value for C4 food; $\sigma$ is a normal distribution of the width; $r_{\mathrm{h}}$ is the radius corresponding to the time of hatching; $c$ is the rate of lens compaction; $r_{\mathrm{T}}$ is the radius corresponding to the start of the C4 feeding period. The residual standard error (SE) is 0.4014 at 362 degrees of freedom. The number of interactions to convergence is 11. The achieved convergence tolerance is $7.463e^{-06}$.

Table S3. Trophic discrimination factors (TDFs) for quail eye lens and primary feathers.

| | Eye lens | | Feather | |
|---|---|---|---|---|
| | $\delta^{13}C$ | $\delta^{15}N$ | $\delta^{13}C$ | $\delta^{15}N$ |
| C3 food | 2.5 | −0.1 | 2.8 | −0.7 |
| C4 food | −0.5 | 1.8 | 0.4 | 1.3 |

Figure S1. Relationship between the lens radius (mm) and δ$^{13}$C values of quail eye lens sections from the control and treatment groups (T10, T15, T20 and T40). The lines were estimated using the piecewise constant function (PCF) model without compaction.

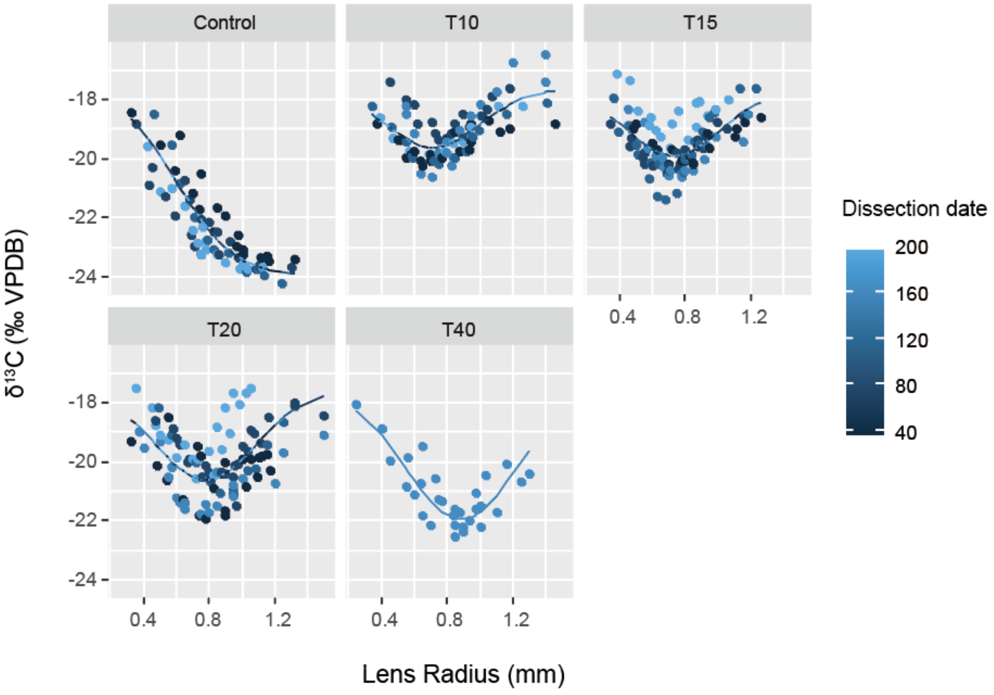

Figure S2. The relationship between the lens radius (mm) and $\delta^{15}N$ values of quail eye lens sections from the control and treatment groups (T10, T15, T20, and T40). The lines were estimated using the piecewise constant function (PCF) model without compaction.

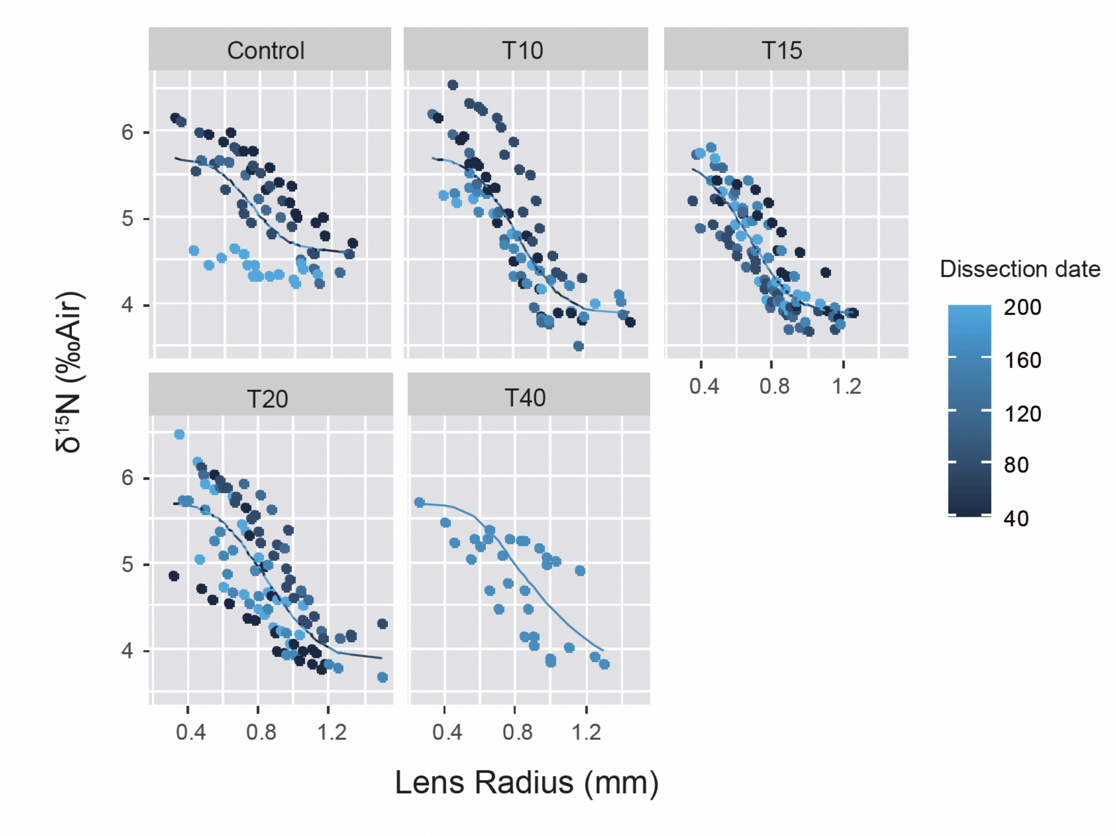

Figure S3. Relationship between the start and end of C3 feeding for each dissection date and transition lens radius ($r_h + r_T$) from the piecewise constant function (PCF) model for $\delta^{13}C$ (a) and $\delta^{15}N$ (b).

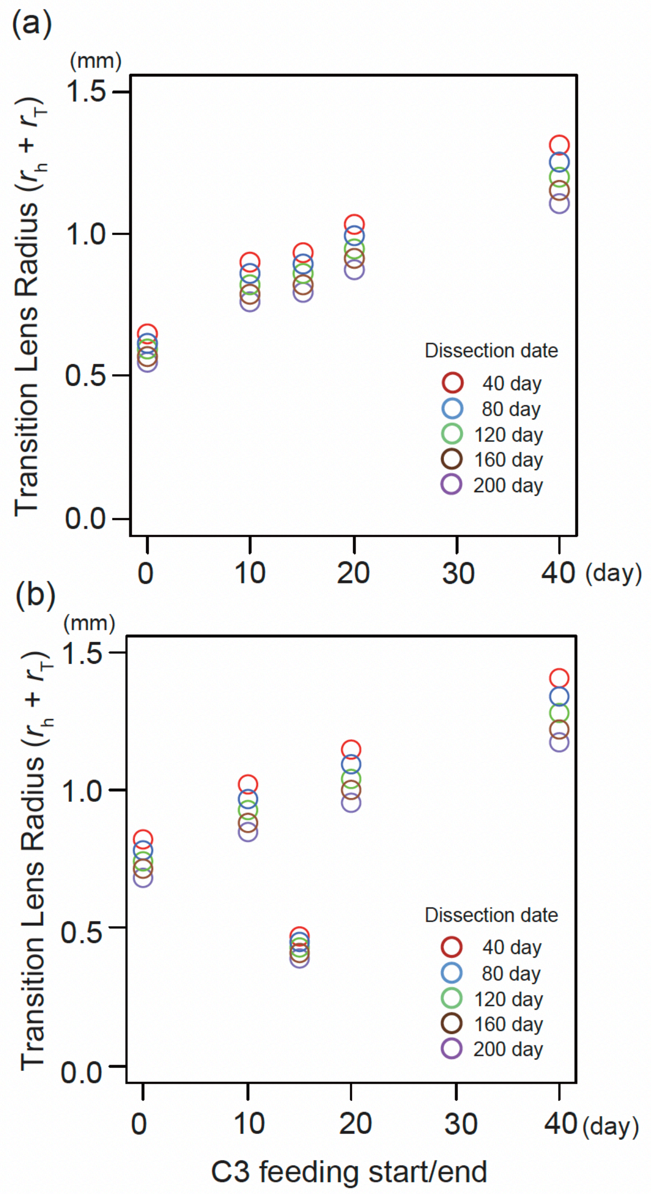

Figure S4. $\delta^{15}N$ values of the quail feather sections: (a) control group (n = 3); (b) T10 group (n = 3); (c) T15 group (n = 3); (d)T20 group (n = 3).

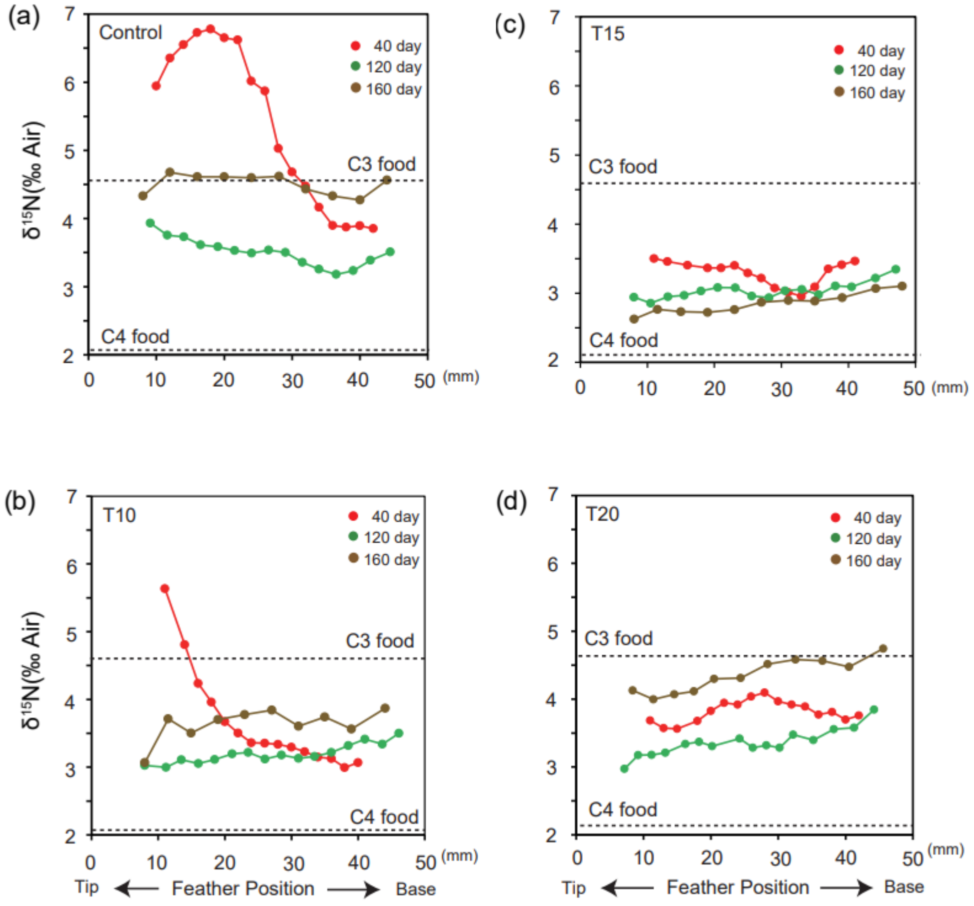
